# Fermentation characteristics of mead and wine generated by yeasts isolated from beehives of two Austrian regions

**DOI:** 10.1101/300780

**Authors:** Helmut Gangl, Ksenija Lopandic, Gabriele Tscheik, Stefan Mandl, Gerhard Leitner, Katharina Wechselberger, Maria Batusic, Wolfgang Tiefenbrunner

## Abstract

Mead is a traditional alcoholic beverage that is produced by fermentation of diluted honey. The mead quality is primarily influenced by the honey variety, although the yeast microflora as the main catalyst of alcoholic fermentation also plays a significant role in the organoleptic and chemical quality of the final product. The impact of the indigenous honey associated-yeasts on the mead properties has scarcely been investigated. To fill this gap the main objective of this work was to assess the metabolic properties of the yeasts isolated from honey and pollen from beehives of northeast Austria.

The biodiversity was low and only two yeast species were identified, *Zygosaccharomyces rouxii* and *Candida apicola.* The fermentation potentials of these yeasts were estimated in two media, grape juice (since yeasts isolated from honey may be useful for sweet wine production) and diluted honey of similar sugar concentration, and compared with those of the reference strains *Saccharomyces cerevisiae*; *S. uvarum* and *S. eubayanus.* Depending on the fermentation substrate, yeasts differed with respect to their metabolic power, fermentation rate, sugar utilization and production of glycerol and organic acids. During mead fermentation *Saccharomyces* species showed the highest metabolic turnover, while the fermentation rate did not differ significantly. Addition of assimilable nitrogen to the diluted honey enhanced fermentation rate of *S. cerevisiae*, but not of the other species. Fermentation of grape juice occurred much faster than that of diluted honey and differences between yeasts were more pronounced. The *S. cerevisiae* commercial wine strain, adapted to high alcohol concentrations, and *S. eubayanus* outperformed the others, *S. uvarum* was comparable with *Z. rouxii*, while *C. apicola* had the lowest fermentation rate. Fructophily of *Z. rouxii* and to a lesser degree of *C. apicola* was observed in both media. An increased production of glycerol was achieved by *S. eubayanus* in both media and by *C. apicola* during the fermentation of honey must. A commercial *S. cerevisiae* strain, *S. eubayanus* and *Z. rouxii* were able to metabolize malic acid in wine. In mead, the *S. eubayanus* and *S. uvarum* yeasts showed the tendency of increasing the level of malic acid. Aroma profile depended profoundly on yeast species. This study demonstrates that the composition and complexity of the fermentation substrate determines the activity and the final metabolic outcomes of the studied yeasts.

## Introduction

Honey is a sweet and flavourful nutrition that is composed mainly of a mixture of carbohydrates and other compounds like amino and other organic acids, proteins, lipids and minerals. Since it is a high sugar, low water medium with low redox potential it causes stress to many microorganisms except to the most osmoresistent, xerophilic specialists (GOMES et al. 2010). Microbial diversity of honey may show regional variation because there are a lot of different contamination sources, such as air, wax feeding organisms, the digestive duct of bees and some that depend on the surrounding of the beehive like nectar and pollen of various flowers and honeydew. Those variation is little investigated although some of the microorganisms, especially yeasts, that survive in or live from honey are of importance either because of their ability to produce fermented beverages or to spoil food products.

Fermented honey, mead, is certainly one of the earliest alcoholic beverages produced by humans. Analyses of ancient pottery jars from the beginning Neolithic (seventh millennium B.C.) in China have revealed that a mix of honey, rice and fruit was used (MCGOVERN et al. 2004). The ingredients of “Nordic Grog” from 1500 B.C. included locally available honey (MCGOVERN et al. 2013). Originally mead was not a pure diluted honey drink (and is not everywhere today). The German name of the dicotyl genus *Philipendula* for instance is a reminder to that; now called “Mädesüß” (sweet girl), it was originally named “Metesüß” (sweet mead; in Norwegian “mjødurt”, mead herb), indicating some importance of the plant for mead aroma enhancement or maybe its flowers served as yeast source.

Mead nowadays is produced using the same purified and cultured yeasts as in wine making (PEREIRA et al. 2013). Historically this was certainly not the case and it’s likely that honey itself was one of the major yeast sources. The objective of this study was to analyze the ability of some indigenous yeasts, directly isolated from beehives of different locations in Austria, to produce mead and sweet wine and to compare their fermentation properties with those of different *Saccharomyces* species.

## Materials and Methods

### • Yeast isolation and selection

Samples were taken from ten beehives from the apiary BeeLocal, located in different landscapes of the northeast of Austria: the wine growing regions Wagram, Weinviertel, Vienna, Neusiedlersee, and the Eastern Alps. Some of these sites (Fig. 1) are in or near a city, others in the proximity of lower mountains or in a xeric grassland beside a steppe lake. Thus the origin of the honey is manifold, from nectar of very different flowers and even honeydew.

**Fig. 1:**
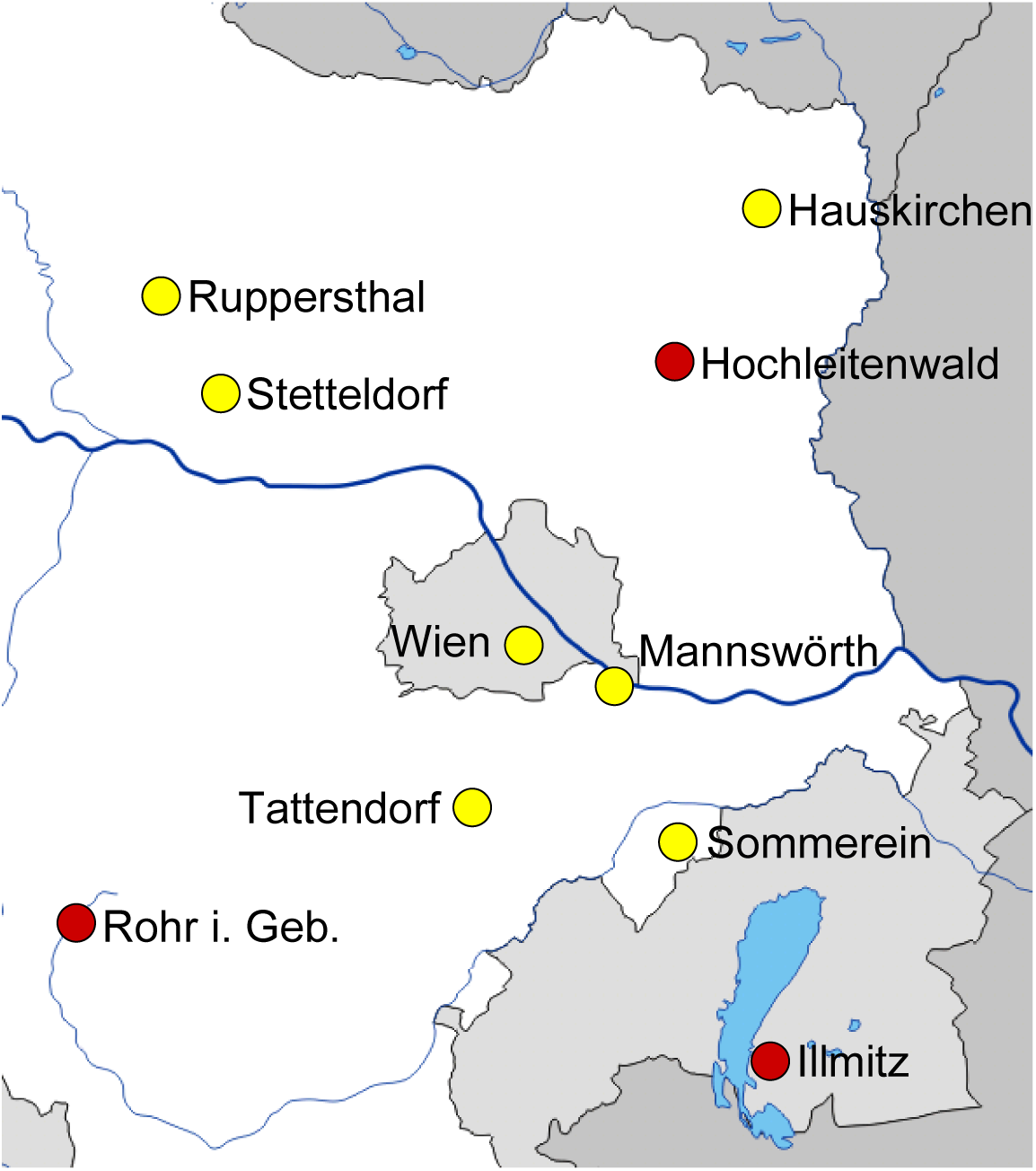
Beehive sample sites in the north-east of Austria. Red circles denote sites where yeast isolation was successful. © d-maps.com.

Samples were taken monthly, from April to beginning of July. Honey and pollen were collected separately out of the honeycombs. All samples were used for yeast isolation in the lab.

The honey and pollen samples were plated on GYP agar (2% glucose, 1% peptone, 0.5% yeast extract). After incubation at 25°C for 3 days colonies were separated and further transmitted until all cells of a culture reached homogeneous appearance. Yeasts were found in the honey samples at the sites Illmitz (surrounded by steppe and hedges), Hochleitenwald (oak forest and arable land) and Rohr im Gebirge (mountainous environment with meadows and pinewood dominated forests). 17 colonies representing three regions were selected and used in genome characterization.

### • Identification of yeast

#### Partial sequencing of the gene coding for 26S rRNA

Yeast DNA was isolated and purified according to the protocol of the MasterPureTM Yeast DNA Purification Kit (Epicentre, Madison, Wisconsin, USA). A fragment of approximately 600 bp of the D1/D2 domain of 26S rRNA encoding gene was amplified in a 50 l reaction mixture containing 1xTris-based reaction buffer pH 8.0 (peqLab, Erlangen, Germany), 4.5 mM MgSO_4_, 0.2 mM of each deoxynucleotide triphosphate, 0.1 pmol l^−1^ of each NL1 and NL4 primer (White et al. 1990), 30 ng of DNA preparation and 0.01 U μl^−1^ of proofreading *Pwo*-DNA-Polymerase (PeqLab, Erlangen, Germany). Polymerase chain reaction (PCR) was performed in a C1000 Thermal Cycler (BioRad, USA) at the following temperature program: 94°C for 2 min of initial denaturation was followed by 35 cycles of 94^o^C for 20s, 54^o^C for 75s and 68^o^C for 80 sec, with a final extension at 68°C for 5 min. The amplicons were purified by Wizard^®^SV Gel and PCR Clean-Up System (Promega, Madison, USA) and sequenced at Microsynth AG (Balgach, Switzerland) with both NL1 (GCATATCAATAAGCGGAGGAAAAG) and NL4 (GGTCCGTGTTTCAAGACGG) primers. The partial sequences of 26S rRNA gene were used in a similarity search by means of the BLAST program from Genbank. The strains showing identity scores higher than 99% were considered conspecific.

### • Microvinification

Four units of water for each one of “Wiener Blütenhonig” (Viennese Flower honey, BeeLocal) were used to produce the diluted medium for mead production. This dilution was chosen to reach sugar concentrations similar to grape juice. Basic chemical parameters are shown in Tab. 1. Inoculation took place at three experiment replications, beginning at the terms 13.05.2015, 14.03.2016 and 18.05.2016. At the latest term, supplementary to the described medium another one was used where initial conditions were changed by adding yeast assimilable nitrogen to the diluted honey so that the content increased from 13 mg/l to 110 mg/l.

**Table 1:**
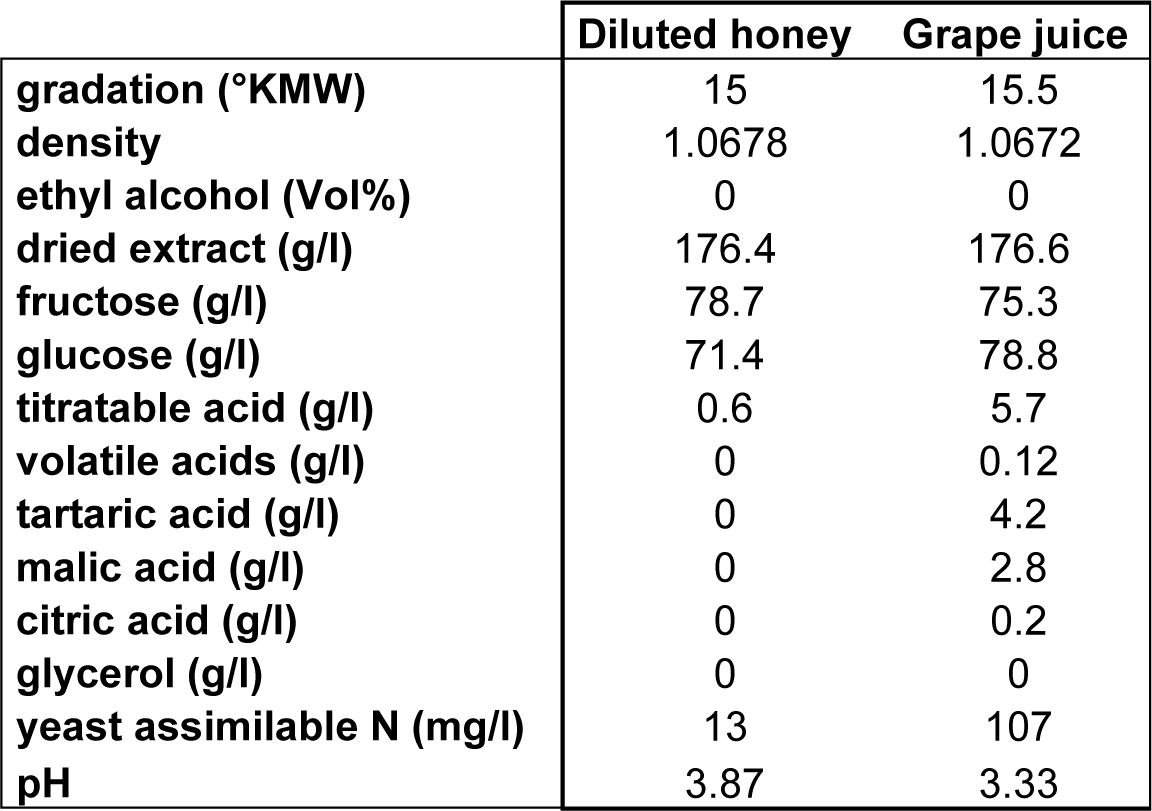
Basic chemical parameters of the original media for fermentation

|  | Diluted honey | Grape juice |
| --- | --- | --- |
| gradation (°KMW) | 15 | 15.5 |
| density | 1.0678 | 1.0672 |
| ethyl alcohol (Vol%) | 0 | 0 |
| dried extract (g/l) | 176.4 | 176.6 |
| fructose (g/l) | 78.7 | 75.3 |
| glucose (g/l) | 71.4 | 78.8 |
| titratable acid (g/l) | 0.6 | 5.7 |
| volatile acids (g/l) | 0 | 0.12 |
| tartaric acid (g/l) | 0 | 4.2 |
| malic acid (g/l) | 0 | 2.8 |
| citric acid (g/l) | 0 | 0.2 |
| glycerol (g/l) | 0 | 0 |
| yeast assimilable N (mg/l) | 13 | 107 |
| pH | 3.87 | 3.33 |

Alternatively a pasteurised grape juice of the Austrian vine variety “Grüner Veltliner” was utilized for wine microvinification beginning on 16.03.2015.

Regardless of the initial medium, yeasts isolated from the locations Hochleitenwald (9 isolates), Illmitz (4 isolates) and Rohr im Gebirge (4 isolates) and reference strains, *S. cerevisiae*, (a laboratory HA 2796^T^ strain and a commercial strain: ‘Pannonia’, Lallemand GmbH, Ottakringer Str. 85-89, 1160 Wien), *S. eubayanus* HA 2841^T^ and *S. uvarum* HA 2786 (isolated from wine must, Tokaj wine region, Hungary) were inoculated in separate flasks. Microvinifications were carried out in 300 ml Erlenmeyer flasks filled with 250 ml diluted honey or grape juice in darkness at a standardised temperature of 20°C. The fermentation progress was monitored by determining the weight loss caused by the production of CO_2_. Fermentation lasted for 23 to 28 days -depending on the initial substrate - before wine chemical analysis started. The resulting wines were tested olfactory. Odour and taste of all wines were acceptable.

### Wine chemical analysis

#### Basic analysis

Chemical analysis of organic compounds of diluted honey, meads, grape juice and wine, like ethyl alcohol and glycerol, sugars (fructose, glucose) and acids (titratable and volatile acids, tartaric, malic, citric and lactic acid) was performed following the OIV Compendium of International Methods of Analysis of Wines and Musts, 2014. Methods for density determination were taken from ALVA-Methodenbuch für Weinanalysen in Österreich, Bundesministerium für Land- und Forstwirtschaft, 1984.

#### Aroma profile of meads

The aroma composition was analyzed, and 63 aroma components from 18 resulting wines were determined. For analysis of volatile aroma compounds the wine sample (8 ml) with 50 μl 3-decanol (48.8 ppm) as internal standard were placed in a 20 ml headspace vial equipped with a magnetic bar and capped with a PTFE-coated silicone septum. For the headspace SPME process a 2-cm DVB/CAR/PDMS (Supelco) fiber Divinylbenzene/Carboxen/Polydimtheylsiloxan, 50/30 μm was chosen to adsorb aroma compounds in gas phase. The fiber was exposed in the headspace of sample vials for 30 min at 30°C.

After the extraction, the fiber was immediately inserted into the GC (GC/MS QP2010, Ultra (Shimadzu)) injection port for 2 min at 250°C for thermal desorption.

For the determination of different aroma compounds a ZB-WAX plus capillary column (length 60 m, 0,5 μm film thickness, 0.25 mm internal diameter; Zebron) was used. The oven temperature was held at 60°C for 3 min before being increased by 10°C/min to 100°C for one minute, and afterwards to 240°C at a rate of 4°C/min and then kept at this temperature for a further 5 min. Helium was used as carrier gas with the constant flow rate of 1.6 ml/min. MS-parameter: Ion Source Temp.: 200°C; Interface Temp.: 250°C; 35-300m/z.

Description of odour and flavour of aroma compounds were taken from http://www.thegoodscentscompany.com/, 07.12.2017.

### • Statistical analysis

A nonlinear regression analysis for the weight loss due to CO_2_ production was performed to determine the fermentation rate. The data were fitted to the sigmoid coursed (HOFBAUER AND SIGMUND 1984) Verhulst equation: y(t) = *γ*/(1+β exp(αt)) which describes the growth of a population when resources are limited. The α parameters are used to estimate the fermentation rate. The coefficient of determination (R^2^), describing how well the data fit to the estimated function, is useful in showing whether the fermentation occurred harmoniously, disturbed or interrupted. Own software was utilized for nonlinear regression. Data had to be prepared to consider initial weight differences and weight loss due to evaporation. In order to assess differences between meads and wines produced by different yeasts, the data of the basic chemical analysis were exported to Statgraphics Centurion Version XV software (StatPoint Inc., Waarenton, VA, USA, 2005) and to ViDaX Version IV (LMS-Data, Trofaiach, Austria, 2005). For mean comparison ANOVA with Levene’s test as variance check and 95% LSD multiple range test were used as well as Kruskal Wallis test and matrix rank permutation test for median comparison. Tests for paired samples were utilized too. Biochemical profiles were evaluated using principal component analysis (Hartung and Elpelt, 1999).

## Results and discussion

### • Identification of yeast isolates

Yeasts used in the mead production practice are usually distinct wine *S. cerevisiae* strains selected in wine and beer fermentation environment (PEREIRA et al. 2008). It is assumed that the yeasts, which are typically adapted to harsh alcoholic fermentation conditions, such as high osmotic pressure, may conduct mead fermentation successfully. However, the chemical composition of grape juice or malted barley is substantially different from that of honey-must, suggesting that these substrates might support the growth of different yeasts (PEREIRA ET AL. 2008, GOMES ET AL. 2010, MENDES-FERREIRA ET AL. 2010). In order to identify the yeast adapted to specific honey properties we isolated 17 yeasts from honey and pollen of beehives at different locations of northeast Austria, which were then used in the fermentation assays of the honey-musts. Comparison of the sequences of the D1/D2 domain of the yeast isolates with those deposited in Genbank suggested two closest relatives *Zygosaccharomyces rouxii* and *Candida apicola.* All yeast isolates (13 strains) from the beehives located in Hochleitenwald (*HLW*-*Zr*) and Illmitz (Illmitz-Zr) were identified as *Z. rouxii*, whereas those four strains from Rohr im Gebirge (*RiG*-*Ca*) belonged to the species *Candida apicola.* Isolation of *Z. rouxii* and *C. apicola* from honey samples is not surprising, as numerous studies have already shown a strong association of these yeast species with high osmotic substrates or bees. *Z. rouxii* is usually isolated from the habitats with increased salt or sugar concentrations, like grape must, wine, jam, wort, salted beans, orange syrup, honey, soft drinks and soy sauce (JAMES AND STRATFORD 2011). On the other hand, the *C. apicola* strains have mostly been isolated from different bee species, although several isolates have also been recovered from pickled cucumbers, grape must and brine in a cheese factory (LACHANCE et al. 2011).

Although the isolated *Z. rouxii* and *C. apicola* are not strong fermenting yeasts like *S. cerevisiae*, a subsequent question arose as to how these indigenous yeasts contribute to the chemical and sensory quality of mead.

### • Fermentation characteristics in mead production

In order to study metabolic power of yeasts during honey fermentation we investigated the basic chemical parameters presented in Tab. 1, which are either products or educts of the fermentation process (e.g. ethyl alcohol and CO_2_ are products, glucose and fructose are educts). Hence these parameters should gather in two correlation clusters. Correlation matrix Fig. 2b highlights significant (significance level α=0.05) correlations between the parameters and shows that on the one hand ethyl alcohol, titratable acids, volatile acids, malic acid and glycerol were positively correlated and on the other hand density, fructose, glucose and pH (most but not all correlations are significant). Between these two groups the correlation was negative. The coherency can be used to compare efficiency of substrate transformations and activity level of yeasts.

**Fig. 2:**
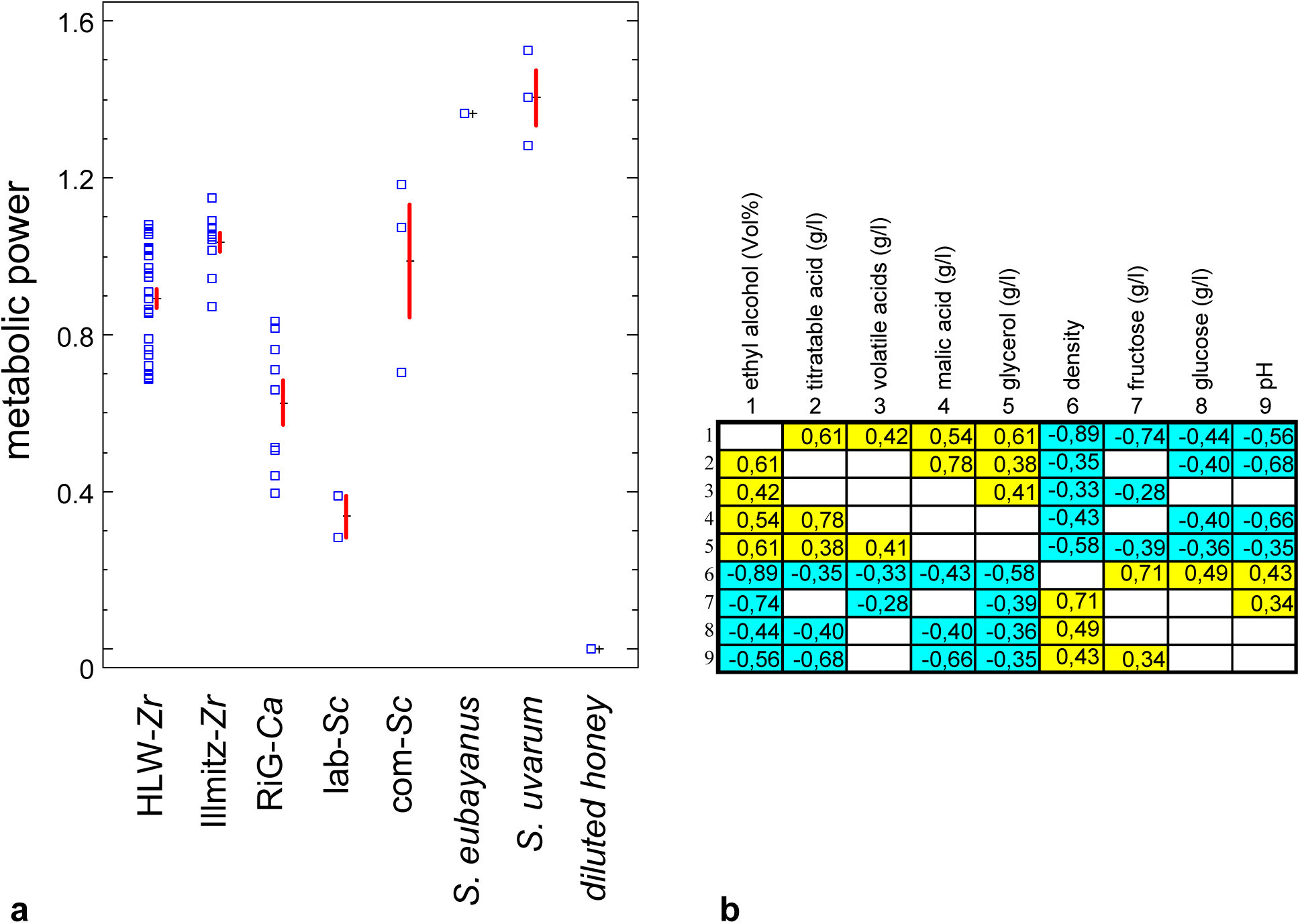
Relative metabolic power (mean and standard error) of the analyzed yeasts (a) and correlation between basic chemical parameters of the meads (b). Yellow: positive correlation. Blue: negative correlation. Significance level α=0.05.

To gain knowledge about relative metabolic power (rMP) of the yeasts the information on the metabolism as well as the correlation between basic chemical parameters were used to create a simple equation:

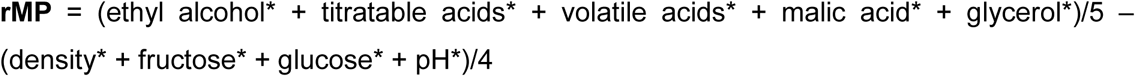

Since the concentrations of the constituents are very different all values are minimum-maximum scaled, so that all parameter values are in the range 0 to 1:

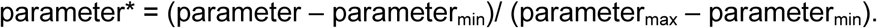

The highest rMPs in mead fermentation were gained by the reference yeasts *S. uvarum* and *S. eubayanus* (Fig. 2a), followed by com-*Sc*, the commercially available pure culture yeast ‘Pannonia’, and Illmitz-*Zr*. The com-*Sc* and Illmitz-*Zr* did not differ significantly (Appendix 1a). The Illmitz yeasts had a significantly higher rMP than the *Z. rouxii* of Hochleitenwald (P=0.001, matrix permutation rank test) despite the fact that they belong to the same species. Yet unknown ecological reasons and selection pressures may cause this difference. *RiG*-*Ca* exhibited significantly less activity in substrate conversion than HLW-*Zr* (P=0.001, matrix permutation rank test) or Illmitz-*Zr* strains (P=0). The lab-*Sc*, a lab yeast not isolated from alcoholic beverages, had the lowest rMP.

In diluted honey fermentation rate (the “speed” of fermentation; coefficients of determination 0.989 – 0.998) did not differ significantly between the yeasts and thus didn’t correlate with metabolic power. On the contrary yeasts differed considerably concerning sugar usage. Since initial glucose concentration was similar to that of fructose, all yeasts below the glucose=fructose line in Fig. 3 are fructophil (more glucose is left) whereas those above are glucophil. Thus all *Saccharomyces* strains used preferably glucose for metabolism. Contrary *C. apicola* utilized slightly more fructose than glucose. This difference is even more pronounced in *Z. rouxii*, which metabolized primarily fructose. This observation is in accordance with the results of PEYNAUD AND DOMERCQ (1955) and LEANDRO et al. (2011). At the level α=0.05 there was no significant disparity of fructose concentration between the HLW-*Zr* and Illmitz-*Zr*-meads (Appendix 1a), but glucose content differed significantly (mean 68.43 g/l HLW-*Zr*-meads versus 64.24 g/l Illmitz-*Zr*-meads). The *RiG*-*Ca*-meads contained on average twice as much fructose than the ones produced by *Z. rouxii*, but the difference concerning glucose concentration was negligible.

**Fig. 3:**
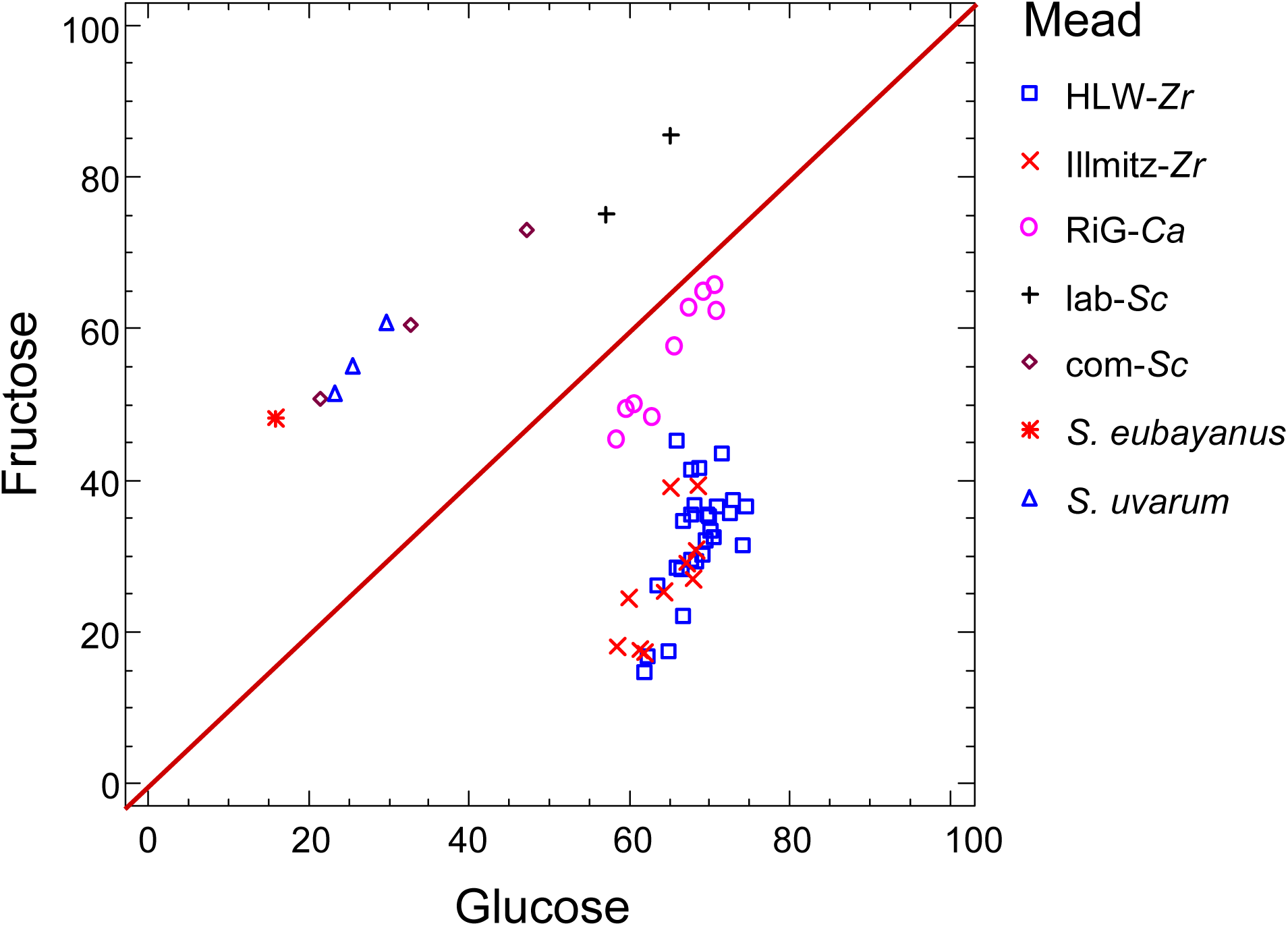
Glucose and fructose content (g/l) of mead.

It has recently been shown that the fructose uptake is related to the presence of different sugar transporters and the ZrFfz1 protein was found to be particularly essential for fructophilic character of *Z. rouxii* (LEANDRO ET AL. 2014). Some *Saccharomyces* species such as *S. eubayanus*, *S. uvarum* and the hybrid yeasts *S. bayanus* and *S. pastorianus* possess in addition to Hxt transporters also a fructose Fsy1 transporter, which catalyzes transport via a proton symport mechanism (RODRIGES DE SOUSA ET AL. 2004, LIBKIND ET AL. 2011, ANJOS ET AL. 2013). The *FSY1* gene is repressed at high sugar concentrations, i.e. the Fsy1 transporter is active only at low fructose concentrations (ANJOS ET AL. 2013, LEANDRO ET AL. 2013), which might explain differences in fructose utilization between *Saccharomyces* yeasts and *Z. rouxii* used in this study.

Ethanol concentrations of meads were generally low and did not exceed 4.8%. Highest concentrations were found in the meads produced by reference yeasts, especially by *S. eubayanus-mead.* Alcohol concentration differed significantly between HLW-*Zr* and Illmitz-*Zr*-meads (Appendix 1a). The ethyl alcohol concentration in *RiG*-*Ca*-meads was on average only half the one of the Illmitz-*Zr*-meads.

There was also a significant distinction between the glycerol concentrations of HLW-*Zr* and Illmitz-*Zr*-meads and between *RiG*-*Ca* and HLW-*Zr*-meads as well. Surprisingly *C. apicola* produced most glycerol (3.01 g/l) of the non-*Saccharomyces* species, but the difference to Illmitz-*Zr* was not significant. Concerning the reference yeasts, *S. eubayanus* was outstanding in glycerol production and *S. uvarum* produced as much as Illmitz-*Zr* and more than both *S. cerevisiae* strains. Glycerol is important for regulating redox potential in the cell and is associated with the cryotolerance in *S. uvarum* and *S. kudriavzevii* (KISHIMOTO ET AL. 1993, PAGET ET AL. 2014). Increased amount of glycerol can significantly contribute to the sweetness and viscosity of wines (NOBLE AND BURSICK 1984). *S uvarum*, *S. kudriavzevii* and their hybrids produce more glycerol than *S. cerevisiae* yeasts in the grape juice fermentations (KISHIMOTO ET AL. 1993, COMBINA ET AL. 2012, PÉREZ-TORADO ET AL. 2015, LOPANDIC ET AL. 2016). *S. kudriavzevii* is able to increase glycerol synthesis by increasing the activity of glycerol-3-phosphate dehydrogenase 1 (*GPD1*) at increased osmotic pressure and low temperature (ARROYO-LÓPEZ ET AL. 2010, OLIVIERA ET AL. 2014). Recently GONZÁLEZ FLORES ET AL. (2017) demonstrated that *S. eubayanus* possess similar properties with *S. uvarum* with respect to the production of glycerol during cider fermentation. This and our study clearly demonstrate that *S. eubayanus* species can ferment and contribute significantly to the viscosity and quality of the mead and other products.

pH-value was highest in *RiG*-*Ca*-*meads* (3.52), indicating low acid production (pH-value of diluted honey was 3.87) and lowest in the ones of *S. uvarum* (2.99), a yeast that produced titratable and malic acids in higher amounts than the others (Appendix 1a). Concentration of titratable acid is low in *RiG*-*Ca*-*meads* (on average 1.53 g/l) and significantly higher in *Z. rouxii* fermented meads, com-*Sc*-mead and *S. eubayanus* one. The difference between these yeasts (with exception of *RiG*-*Ca*) was not significant but *S. uvarum*-mead – which had, as already mentioned, the highest values (4.23 g/l on average; Appendix 1a) – differed significantly from them. Volatile acids concentrations did not differ significantly between the meads (ANOVA P=0.53; Kruskal Wallis P=0.52). Substantial differences were observed in the production of malic acid (Appendix 1a). Malic acid content was zero in *RiG*-*Ca* and *S. cerevisiae* (both strains), significantly higher in *Z. rouxii* and the highest concentrations were observed in the meads generated by *S. eubayanus* (1.1 g/l) and *S. uvarum* (1.6 g/l). Significant differences are denoted in Appendix 1a. It was already shown that *S. uvarum* has been able to synthesize malic acid during wine fermentation (RAINERI AND PRETORIUS 2000). This trait is very important for the wine production in areas with hot climates, where grape juice acidity is usually low (CASTELLARI ET AL. 1994). Results of our study have shown that beside *S. uvarum*, *S. eubayanus* can also produce increased concentration of malic acid during mead fermentation (Appendix 1a).

Lactic and citric acid content was low in all meads. Surprisingly meads created by *Z. rouxii* tasted sourer than the ones fermented with the aid of *Saccharomyces* species.

### • Artificial nitrogen addition

The available nitrogen concentration for the yeasts in diluted honey was only about one eighth of that in the grape juice. Since shortage of nitrogen may slow down or even stop fermentation, in one experiment assimilable nitrogen was added to the diluted honey. The t-test and signed rank test for paired samples was performed to analyze the influence of additional nitrogen (N+). Some of the results are counterintuitive.

If all meads are considered, no matter which yeast species produced them, on average fructose concentration was significantly higher in the N+ meads and thus was less utilized (P=0.03, t-test). However, this was only the case in HLW-*Zr* and Illmitz-*Zr*-meads, not in RiG-Ca-meads and especially not in com-*Sc*-mead: ‘Pannonia’ did much better with additional nitrogen (N+ mead: 38.2 g/l fructose; N-mead: 60.5 g/l). Concerning this yeast strain, the difference in glucose concentration was even higher (7.3 g/l versus 32.6 g/l) but on average yeasts used glucose equally in both experimental variants (P=0.054, t-test).

Without exception all yeasts produced more glycerol in the N+ mead variant (P=0.0 t-test), but not more ethyl alcohol (P=0.083, signed rank test).

Tartaric acid was detected in significant higher amount (P=0.0004 signed rank test) in the N+ meads, whereas the concentrations of malic acid were usually lower (P=0.001 signed rank test) in the N+ meads. However, *RiG*-*Ca* did not produce malic acid. The pH-value was higher in the N+ variant (P=0.0007 signed rank test), indicating that more acids were produced in the N-meads in the whole.

Besides, there was a very distinct influence on fermentation rate (Fig. 4). Nitrogen addition increased fermentation in *S. cerevisiae*, but had no impact on rate of fermentation of *C. apicola*, and decreased metabolic speed in *Z. rouxii* and the *S. uvarum* reference strain.

**Fig. 4:**
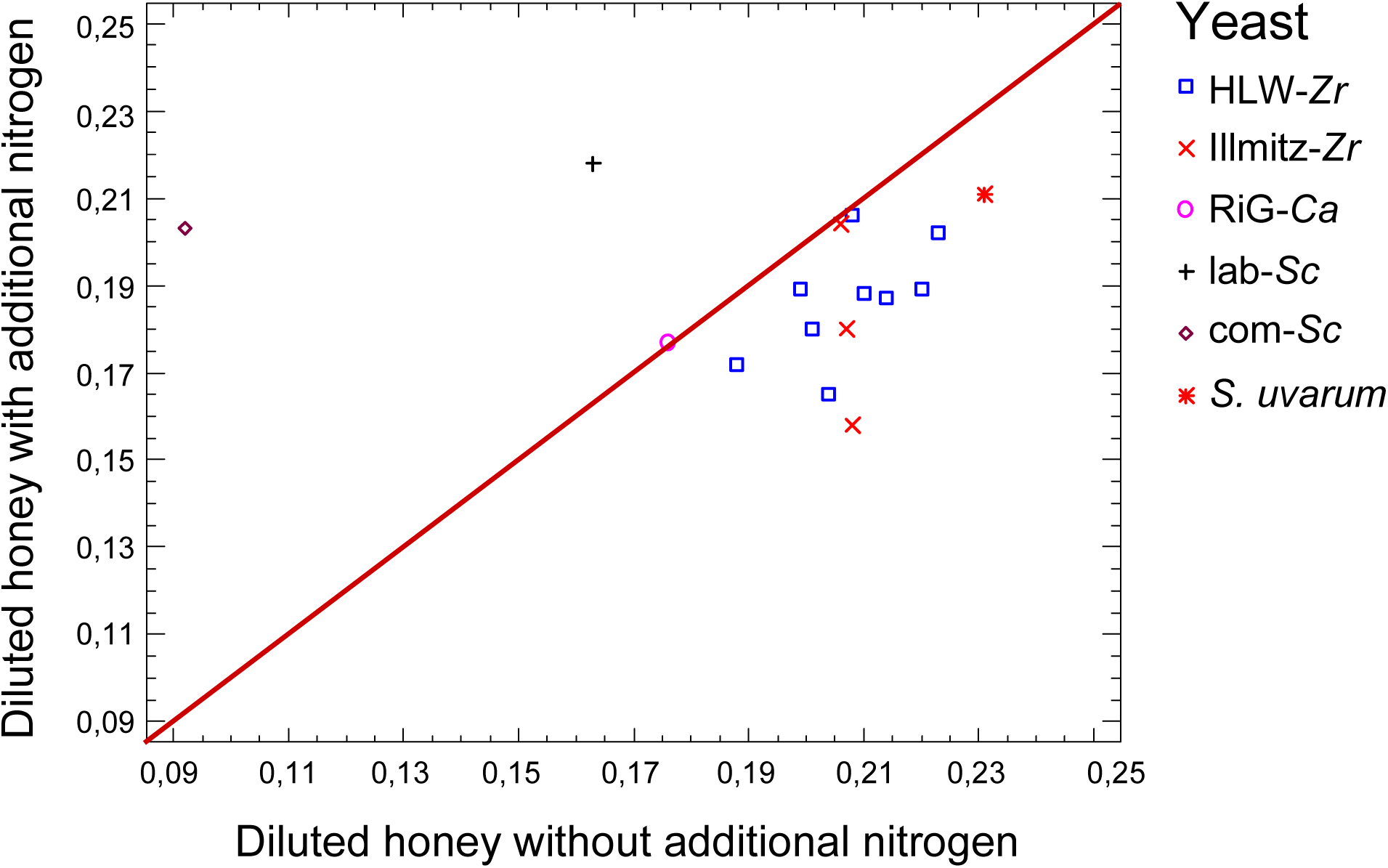
Logarithm of fermentation rate during mead production with and without additional assimilable nitrogen. Comparison of meads produced by different fermenting yeasts.

### • Comparison of the fermentation characteristics of mead and wine

Whereas fermentation rates of all yeasts were very similar during mead production, there were huge rate differences in the process of wine making. The com-*Sc* ‘Pannonia’ and the *S. eubayanus* reference strain had highest fermentation rates and RiG-*Ca* the lowest. Nevertheless *C. apicola* fermentation of grape juice was faster than any fermentation of diluted honey whatever yeast was used (Fig. 5).

**Fig. 5:**
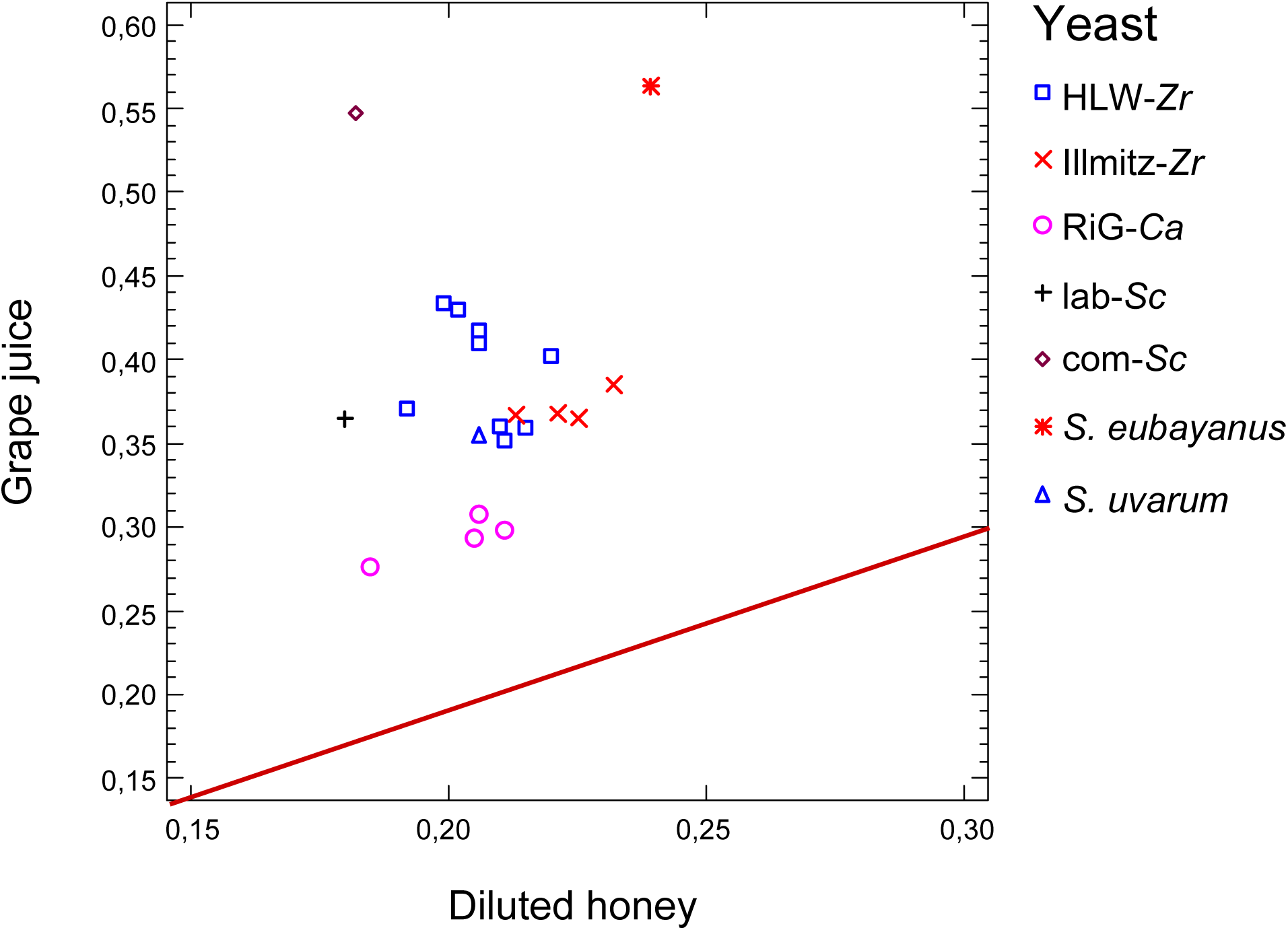
Logarithm of fermentation rates in mead and wine production. Comparison of meads and wines produced by different fermenting yeasts. The red line indicates an identical fermentation rate in mead and wine production. Note the scale difference.

Like in diluted honey a preferred use of one of the monosaccharides glucose and fructose in wine production was observed (Appendix 1b). In the wine generated by *S. uvarum* and lab-*Sc* there was more fructose left than glucose. Differently from the honey fermentation, the com-*Sc* and *S. eubayanus* consumed merely all of both monosaccharides during the wine fermentation. The fructophilic nature of the *Z rouxii* and *C. apicola* was also characteristic for the wine fermentation. *C. apicola* consumed less sugar than *Z. rouxii.* Altogether with exception of *S. uvarum* and lab-*Sc*, sugar consumption of yeasts was comparable in both media. In the grape juice concentration of both sugars was initially similar (Tab. 1).

Alcohol concentration was up to 9% in wines fermented by *S. eubayanus* and com-*Sc*, much higher than in meads (4.8%). As expected, *Z. rouxii* and *C. apicola* produced much less ethanol, 1.34% and 1.89% respectively (Appendix 1a and 1b).

In winemaking, the concentration of titratable acid is a quality indicator, since a too high or too low pH can alter organoleptic properties of wine. All yeasts produced much more titratable acids during wine than mead production. In both media *S. uvarum* was outstanding (8.15 g/l in wine and 4.23 g/l in mead) and differed significantly in the production of titratable acids from the other yeasts (ANOVA P=0.025 and Multiple Range Test). Concentration of volatile acids were in the range of 0.22 g/l to 0.39 g/l. There was no significant difference between the wines. Tartaric acid content was very different in honey and grape juice and therefore also in mead and wine. Concentration in wines was about 4 g/l (Appendix 1b). The content of malic acid in honey was zero like that of tartaric acid. Malic acid concentrations comparable with that of grape juice (2.8 g/l) existed in RiG-*Ca*-wine, lab-*Sc*-wine and *S. uvarum*-wine, whereas the content was significantly lower in *Z. rouxii* fermented wines and in *S. eubayanus-wine* and com-*Sc*-wine. Obviously *Z. rouxii*, *S. eubayanus* and *S. cerevisiae* commercial strain ‘Pannonia’ are able to metabolize malic acid. In winemaking, malic acid contributes particularly to the acidic taste and may support the growth of lactic acid bacteria leading to wine spoilage after bottling (PRETORIUS 2000, REDZEPOVIC et al. 2003). Therefore, the *Saccharomyces* strains that are able to utilize malic acid are highly desirable in the wine production, especially in the area with cold climate, where the acidity of grape juice is usually increased (RAINIERI AND PRETORIUS 2000). It was reported that thermotolerant *Saccharomyces* yeasts may convert up to 50% of malic acid into ethanol, while the cryotolerant strains synthesize malic acid rather than degrade (CASTELLARI ET AL. 1998). However, the majority of *S. cerevisiae* cannot efficiently degrade L-malic acid during alcoholic fermentation (for review see VOLSCHENK ET AL. 2003). The inefficient degradation of L-malic acid by *S. cerevisiae* is ascribed to the slow uptake of L-malic acid by diffusion and the low substrate affinity of its malic enzyme (Km=50 mM) located in mitochondria (BARANOWSKI AND RADLER 1984, SALMON 1987, VOLSCHENK ET AL. 1997). Our results also show that the ability of yeasts to utilize malic acid is not only species and strain-dependent, but the growth substrate may also play a significant role. As already mentioned, the cryotolerant *S. eubayanus* showed different behavior regarding malic acid utilization in mead and grape juice environment.

*Zygosaccharomyces bailii* has been described as the yeast which can metabolize high concentration of malic acid (BARANOWSKI AND RADLER 1984). In our study, another species *Z. rouxii* was also shown to be able to utilize malic acid in must fermentation, and the production of low concentration of malic acid in mead fermentation was also observed (Appendix 1a). Just to mention that the commercial production of malic and succinic acids by *Z. rouxii* at 30% glucose was already studied (TAING AND TAING 2007). Based on the results of our study the potential application of *S. eubayanus* and *S. uvarum* in the production of malic acids could also be considered (Appendix 1a).

Citric acid concentrations of wines were about the same as in the initial grape juice. Lactic acid concentrations were much higher in com-*Sc*-wine and *S. eubayanus*-*wine* and comparable low in all others. Glycerol concentrations in wines were higher (on average 3.31 g/l) than in meads (mean: 2.56 g/l), especially in the one of com-*Sc* (4.5 g/l). No significant difference between wines fermented by *Z. rouxii* (3.3 g/l HLW-*Zr*; 3.5 g/l Illmitz-*Zr*) and *C. apicola* (3.2 g/l on average) was detected.

pH-value of grape juice (3.33) was lower than the one of diluted honey (3.87) whereas the pH of wines (3.27) and meads (3.3) were similar. Wines fermented by C. apicola, com-*Sc* and S. uvarum had pH values 3.41, 3.21 and 3.13, respectively. Within *Z. rouxii* there was a significant difference (Multiple Range Test): HLW-*Zr*-wines had a lower pH than the Ilmitz-*Zr*-wines.

In wine production the utilization of assimilable nitrogen was low in *C. apicola* and lab-*Sc* and very high in com-*Sc*. *Z. rouxii*, *S. uvarum* and *S. eubayanus* laid in the same range.

## Mead aroma composition

Concerning odour, a quantitative description of aroma profile may not be very informative since olfactory perception is not additive. Up to now there is no generally accepted theory of olfaction. Molecular shape, volume and vibration in the IR realm are recognized (FRANCO et al. 2011, SABERI & SEYED-ALLAEI 2016, PAOLI ET AL. 2016) and the simultaneous sensations of different molecules are superimposed in a yet unknown way to an overall impression. The physical principles of sensation are quite different from the ones of technical measurement and identification of aroma compounds. Although all (or most) humans perceive the same at receptor level, the interpretation of the signal in the brain may be very different and thus subjective. Nevertheless the main reason for chemical analyses of aroma compounds is analytical support of sensorial evaluation and thus we try to meet some conclusions.

The basic smell of wine is produced by by-products of alcoholic fermentation like methanol, ethanol, fusel alcohols, acids and their esters. These compounds in the right concentration distinguish the typical flavour of wine from other aromas, they create the basic odour of all wines, no matter of red or white and independent of variety. It even differs from the basic odour of the must and thus is likely important to separate typical wine from mead smell, too.

Beyond that, wine flavour contains many more components, like terpenes, phenols, aldehydes, etc. Although existing in minor quantity they are intensive in smell and create the peculiarity of wine varieties. In most varieties only a few aroma compounds are responsible for typical flavour (e.g. linalool for Muskat, 1,8-cineol for currant smell, iso-amylacetate for banana odour is characteristic of a typical Welschriesling). These compounds may be more important to characterize in meads made by different yeast species and strains.

To distinguish mead from wine, the aroma profile of meads were compared with those of a white wine (Weißer Storch, Lenz Moser AG, Krems, Austria), with a typical harmonic, neutral and unobtrusive flavour (Tab. 2 and Tab. 3). Compared to grape wine aroma profile of mead contains more terpenes (Tab. 2) which are also characteristic of essential oils (e.g. borneol, thymol, rose-oxide, nerol oxide, etc.) produced by a variety of plants. Compounds with honey, sweet, creamy, camphor balsamic, floral, rose, spicy and woody odours dominate mead aroma. They are atypical for all grape wines, but some of these compounds are also contained in wines with a special aroma resulting from high levels of terpenes (e.g. Muskat, Traminer, etc.).

**Table 2:** Aroma compounds that are typical for mead. Relative concentration in percent.

| compound | white wine | mead n=18 |
| --- | --- | --- |
| Ethylpentadecanoate | 0 | 100 |
| Acetoin | 0 | 100 |
| iso-Amyldecanoate | 0 | 100 |
| Borneol | 0 | 100 |
| Propanoic acid, 2-phenylethyl ester | 1 | 99 |
| Thymol | 1 | 99 |
| p-Cymen-8-ol | 1 | 99 |
| p-Cymenene | 1 | 99 |
| rose oxide -trans | 2 | 98 |
| Eugenol | 2 | 98 |
| Nonanal | 3 | 97 |
| Ethylnonanoate | 3 | 97 |
| Ethyllaurate | 3 | 97 |
| 1-Butanol | 3 | 97 |
| alpha-Terpinene | 4 | 96 |
| rose oxide -cis | 4 | 96 |
| Hotrienol | 6 | 94 |
| 3-Decanone | 8 | 92 |
| Terpinen-4-ol (Carvomenthenol) | 10 | 90 |
| p-Cymene | 12 | 88 |
| 1-Nonanol | 12 | 88 |
| Decanal | 15 | 85 |
| 2-Nonanol | 15 | 85 |
| Ethylpalmitate | 16 | 84 |
| Amylpropionate | 18 | 82 |
| Ethyl-iso-valerate | 18 | 82 |
| Nerol oxide | 19 | 81 |
| alpha-Phellandrene | 20 | 80 |

Relative to mead, in white wine very fruity, green and flowery odours dominate (Tab. 3).

**Table 3:**
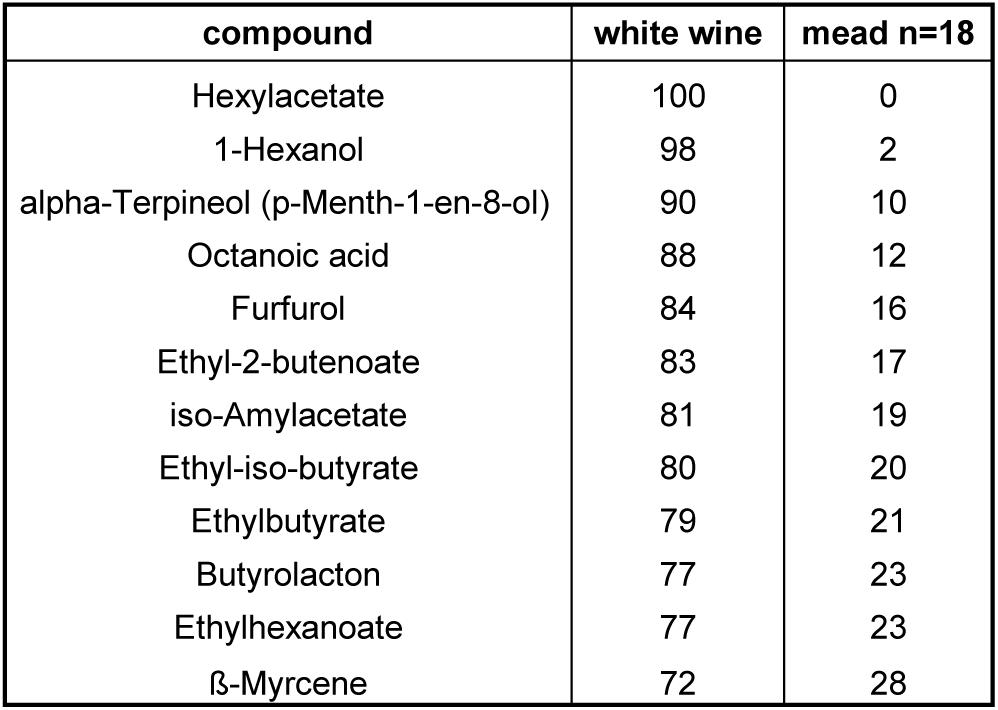
Aroma compounds that are typical for wine compared with mead. Relative concentration in percent.

Aroma profile depends on fermenting yeast (SWIEGERS ET AL. 2005, STYGER ET AL. 2011). Most typical for *Z. rouxii* wines was 2-butenoic acid, ethyl ester, which was at least five times as concentrated as in other wines. However, in Illmitz-*Zr* wines it was more than double as concentrated as in HLW-*Zr* ones. The smell is pungent, chemical diffusive, sweet, alliaceous, like caramel or rum. Typically, very intensive honey and flowery, rose, sweet smell results from 2-phenylethylacetate and -propionate. 2-phenylethylacetate is a basic compound in grape wine and was equally concentrated in *S. uvarum* wine as in *Z. rouxii* mead, but not in *S. cerevisiae* or *C. apicola* ones. Contrary, *Z. rouxii* was the sole producer of higher amounts of 2-phenylethylpropionate, which is not a basic compound of grape wine. Furthermore the fusel alcohols isobutanol and 2-nonanol were more concentrated in *Z. rouxii*-than in other meads. Important odour components like eugenol (intensive sweet, spicy, clove, woody) and ethylhexanoate (fruity) were lower concentrated in *Z. rouxii* meads than in others.

*Z. rouxii* is most important in producing the flavour molecules of soy sauce, the main aroma compounds are 4-hyroxy-3[2H]-furanones with attractive flavour and low odour thresholds, especially 4-hyroxy-2,5-dimethyl-3[2H]-furanone (HAUCK et al. 2003). However, we could not detect this substance in *Z. rouxii* produced meads or others.

In *C. apicola* meads furfurol with sweet, woody, almond, fragrant, baked bread odour, dominates, linalool oxide concentration was high (floral, sweet, woody) compared with other meads, whereas rose oxide had the lowest concentration.

The *S. uvarum*-mead was distinct by high hexylacetate concentration with an intensive, fruity, green, apple, banana and sweet odour. Rose oxid with rosy, green, floral, spicy smell and amyloctanoate and –decanoate with waxy, oily, fruity, green and cognac flavour were also typical for this very aromatic mead.

The aroma profile of Com-*Sc* mead was more like the one of *S. uvarum* but with a lower level of important compounds, e.g. isobutyl-, hexyl- and 2-phenylethylacetate (fruity, flowery, tropical, banana, sweet odour). Linalool oxide (earthy, floral, sweet, woody) concentration was raised.

PCA confirms that aroma composition of meads depends on the species that produced them. Principal component 1 (PC1) very clearly separates the *Z. rouxii* mead aromas from all others, whereas the aroma compositions of HLW-*Zr* and Illmitz-*Zr*-meads did not show any remarkable differences (with the already mentioned exception of 2-butenoic acid, ethyl ester concentration) and thus origin of yeast strains is negligible concerning production of aroma compounds. This is not self-evident since rMP and concentration of glucose, ethyl alcolhol and glycerol differed significantly between HLW-*Zr*-meads and Illmitz-*Zr*-meads (Appendix 1a).

PC2 orders meads in roughly the same way as rMP, although in opposite direction: lowest values are reached by *S. uvarum*-meads and highest in *C. apicola* ones. This indicates that high metabolism in general is the key for elevated aroma compound production. Indistinguishable are the PC1-values of com-*Sc*-mead and meads produced by *Z. rouxii.*

**Fig. 6:**
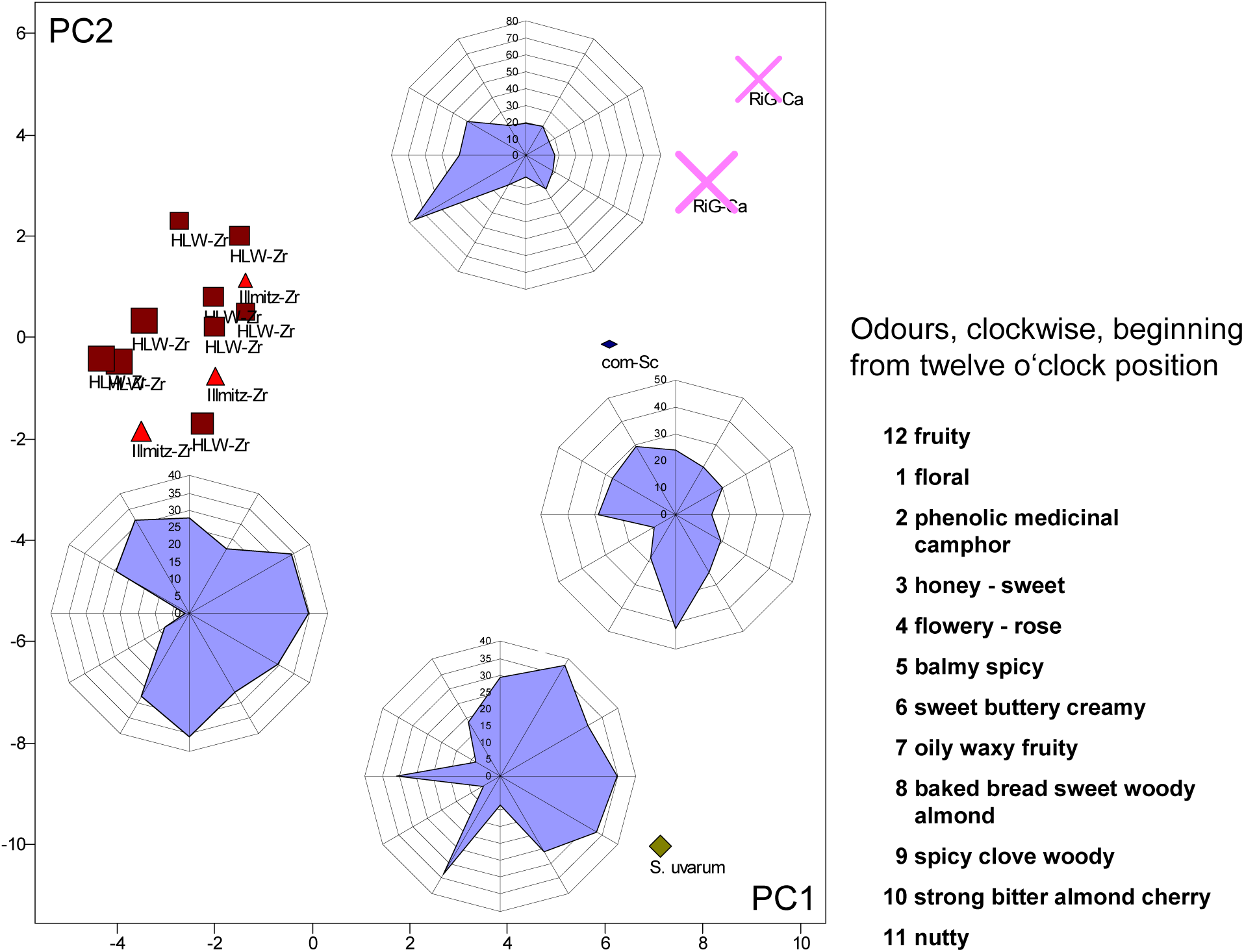
Principal component analysis of mead aroma profiles and circular charts for odour description (see also Appendix 2).

On the whole, every yeast species creates its own, characteristic mead flavour and odour; especially *Z. rouxii* meads are certainly an interesting alternative to those produced by *Saccharomyces* yeasts. Their aroma may be more in accordance with the mead smell and taste from the past when no commercially utilized yeasts were available.

## Conclusions

The results of the present study have shown remarkable differences in the fermentation and aroma performances of the yeasts used for the mead and wine production. As expected, yeasts of the genus *Saccharomyces* showed the greatest efficiency in metabolic turnover, especially *S. uvarum* and *S. eubayanus.* In mead fermentation, *S. cerevisiae* was more inhibited by low level of assimilable nitrogen than all other yeasts that didn’t benefit from addition of nitrogen. Nevertheless a proper strain selection here might generate the product of preferred quality.

Immediate alcohol production is associated with high fermentation rate and guarantees low risk of unwanted microorganisms inhabiting the medium during early fermentation. Since this early phase is especially important for aroma component creation high fermentation rates are significant for taste and odour of the product. However, contrary to wine fermentations in mead production yeasts of the genus *Saccharomyces* did not show significantly higher fermentation rates than *Z. rouxii* and *C. apicola*, respectively. From this point of view all yeasts were equally appropriate.

To our knowledge, it was the first time that *S. eubayanus* was used in the fermentation of the high-sugar mead and grape juice and it showed valuable properties in comparison to the other yeasts. *S. eubayanus* is a cold-tolerant yeast, which in the association with *S. cerevisiae* generate the interspecies hybrids known as *S. pastorianus* or lager brewing yeasts (Libkind et al. 2011). Fermentative ability of *S. eubayanus* have scarcely been studied to date and a recent study has shown that this yeast exhibits low fermentation performance of malt worth and inability to use maltotriose (GIBSON et al. 2013). GONZÁLEZ FLORES ET AL. (2017) also demonstrated that *S. uvarum* and *S. eubayanus* possess similar properties regarding the production of increased amount of glycerol, low acetic acid and elevated production of the 2-phenylethanol and 2-phenylethyl acetate during cider fermentation. Our results also show that *S. eubayanus* is able to generate a high level of glycerol during mead and wine production. Glycerol contributes to the viscosity, sweetness and complexity of flavour and makes the taste of the final product creamier suggesting that *S. eubayanus* could improve organoleptic properties of the final products. In this regard it is worth mentioning that for all yeasts nitrogen addition increased glycerol production during diluted honey fermentation and thus may enhance mead quality although it inhibited fermentation rate of most yeasts.

Fructophily in yeasts may reach considerable practical importance since it has turned out recently that there is a relationship between high intake of fructose – sweetened beverages and type 2 diabetes. Surprisingly only fructose, but not glucose, impairs insulin signalling (BAENA et al. 2016). Furthermore fructose intake enhances gout risk (CHOI et al. 2010). The fructophilic yeast *Z. rouxii* therefore may gain additional importance in the production of alcoholic beverages such as mead and winen. In a possible scenario *Z. rouxii* may be used at the beginning of fermentation and afterwards more alcohol tolerant yeasts that ferment glucose more rapidly than fructose and that may even benefit from the low fructose level. However, further research is needed to prove the usefulness of mixed or successive approaches (JOLLY et al. 2014).

## Acknowledgements

We thank Astrid Tiefenbrunner for critical remarks.

---

**Appendix 1a:**
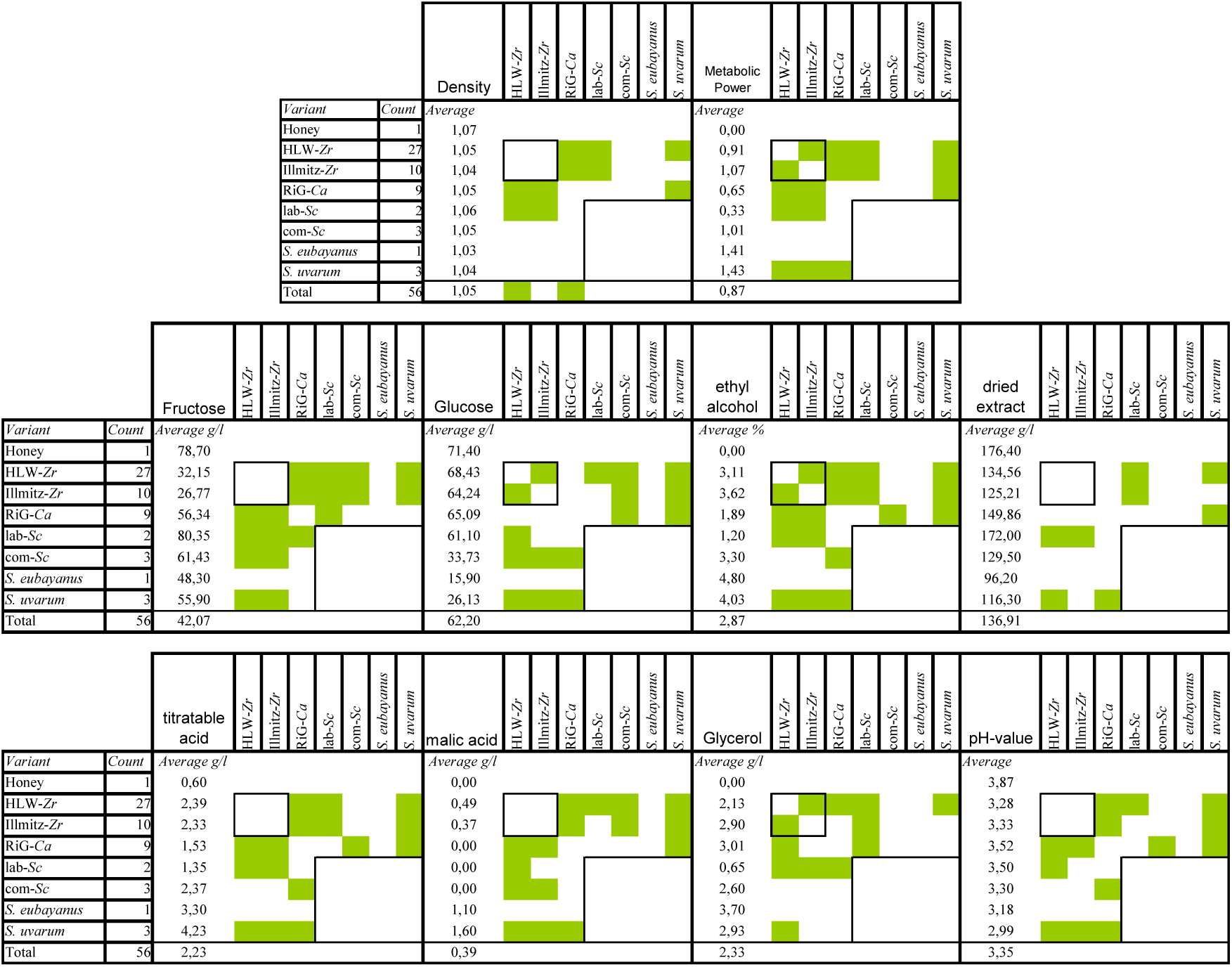
Significant difference (significance level α=0,05; P<α) between basic chemical parameters in mead variants.

**Appendix 1b:**
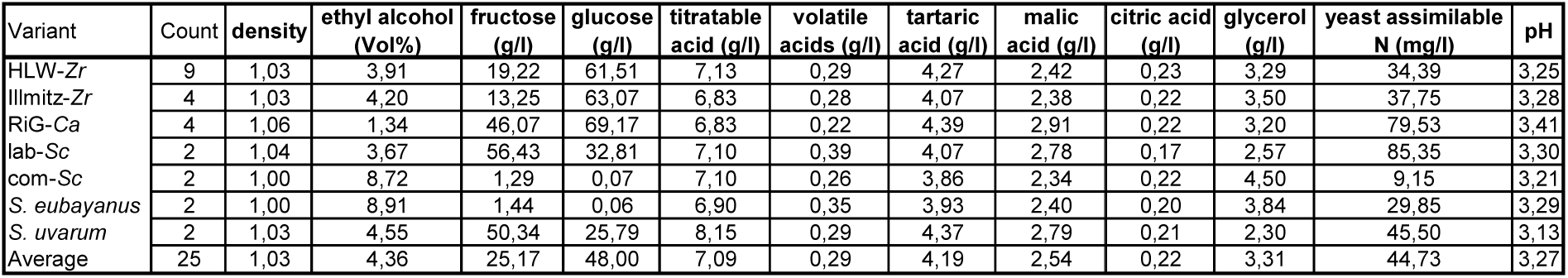
Basic chemical parameters in grape wine variants

**Appendix 2:**
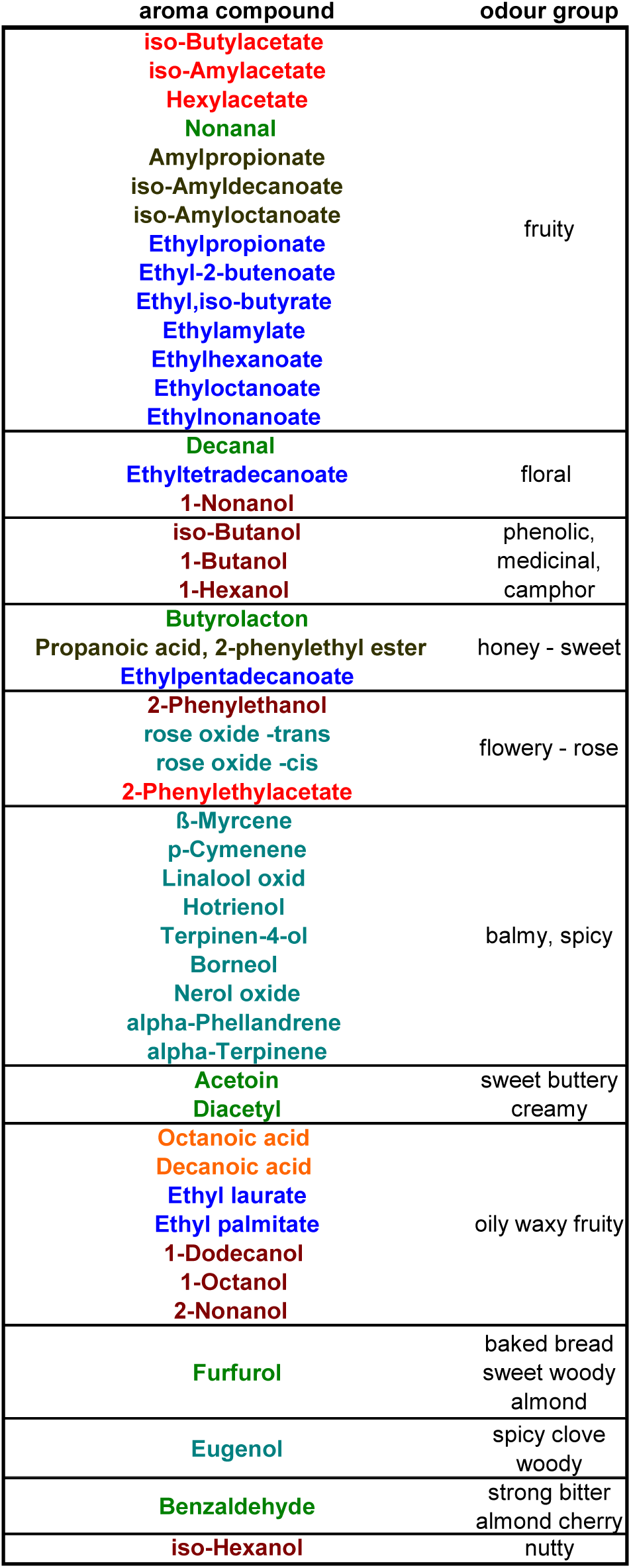
Odour specification for the circular charts of Fig. 6

## References

ALVA-Methodenbuch für Weinanalysen in Österreich, Bundesministerium für Land- und Forstwirtschaft, 1984.

Anjos, J., Rodriges de Sousa, H., Roca, Ch., Cássio, F., Luttik, M., Pronk, J.T., Salema-Oom, M., Gonçalves, P., 2013. Fsy1, the sole hexose-proton transporter characterized in *Saccharomyces* yeasts, exhibits a variable fructose: H^+^ stoichiometry. Biochim Biophis Acta, 1828, 201–207.

Arroyo-López, F.N., Orlic, S., Querol, A., Barrio, E., 2009. Effects of temperature, pH and sugar concentration on the growth parameters of *Saccharomyces cerevisiae, S. kudriavzevii* and their interspecific hybrids. Int J Food Microbiol 131, 1120–127.

Baena, M., Sangüesa, G., Davalos, A., Latasa, M-J., Sala-Vila, A., Sanchez, R.M., Roglans, N., Laguna, J.C., Alegret, M., 2016. Fructose, but not glucose, impairs insulin signaling in the three major insulin-sensitive tissues. Scientific Reports, 6, 26149, 1–15.

Baranowski K., Radler F., 1984. The glucose-dependent transport of L-malate in *Zygosaccharomyces bailii.* Antonie Van Leeuwenhoek 50, 329–340.

Castellari L., Tini V., Zambonelli C., Rainieri S., 1998. Oenological properties of cryotolerant and thermotolerant *Saccharomyces* strains. Food Technology and Biotechnology 36, 319–323.

Choi, H.k., Willet, W., Curhan, G., 2010. Fructose-Rich Beverages and Risk of Gout in Women. Journal of the American Medical Association, 304(20), 2270–2278. doi:10.1001/jama.2010.1638.

Combina, M., Perez-Torrado, R., Tronchoni, J., Belloch, C., Querol, A., 2012. Genome-wide gene expression of natural hybrid between *Saccharomyces cerevisiae* and S. *kudriavzevii* under enological conditions. Int J Food Microbiol 157, 340–345.

Franco, M.I., Turin, L., Mershin, A., Skoulakis, E.M.C., 2011. Molecular vibration-sensing component in *Drososphila melanogaster* olfaction. Proc. Natl. Acad. Sci. USA. 108(9), 3797–802. doi:10.1073/pnas.1012293108. Epub 2011 Feb 14.

Gangl H., Batusic M., Tscheik G., Tiefenbrunner W., Hack C., Lopandic K., 2009. Exceptional fermentation characteristics of natural hybrids from *Saccharomyces cerevisiae* and *S. kudriavzevii*. New Biotechnol 25, 244–251.

González Flores M., Rodrígez M.W., Oteiza J.M., Barbagelata R.J., Lopes C.A., 2017. Physiological characterization of *Saccharomyces uvarum* and *Saccharomyces eubayanus* from Patagonia and their potential for cidermaking. Int. J. Food. Microbiol. 249, 9–17.

Gibson, B.R., Storgards, E., Krogerus, K., Vidgren, V., 2013. Comparative physiology and fermentation performance of Saaz and Frohberg lager yeast strains and the parental species *Saccharomyces eubayanus*. Yeast 30, 255–266.

Gomes, S., Dias, L.G., Moreira, L.L., Rodrigues, P., Estevinho, L., 2010. Physicochemical, microbiological and antimicrobial properties of commercial honeys from Portugal. Food and Chemical Toxicology 48, 544–548.

González, S.S., Gallo, L., Climent, M.D., Barrio, E., Querol, A., 2007. Enological characterization of natural hybrids from *Saccharomyces cerevisiae* and *Saccharomyces kudriavzevii*. Int. J. Food Microbiol. 116, 11–18.

Hartung, J., Elpelt, B., 1999. Multivariate Statistik, Oldenburg Wissenschaftsverlag.

Hofbauer, J., Sigmund, K., 1984. Evolutionstheorie und dynamische Systeme. Mathematische Aspekte der Selektion. Parey, Berlin-Hamburg.

James S.A., Stratford M., 2011. *Zygosaccharomyces* Barker (1901). *In* The Yeasts, a Taxonomic Study, 5th edn, Vol. 2, ed. by Kurtzman, C.P., Fell, J.W. and Boekhout, T., Elsevier, London, pp. 937–947.

Jolly, N.P., Valera, C., Pretorius, I.S., 2014. Not your ordinary yeast: non-*Saccharomyces* yeasts in wine production uncovered. FEMS Yeast Res. 14, 215–237.

Kishimoto, M., Goto, S., 1995. Growth temperatures and electrophoretic karyotyping as tools for practical discrimination of *Saccharomyces bayanus* and *Saccharomyces cerevisiae*. J. Gen. Appl. Microbiol. 41, 239–247.

Lachance M.A., Boekhout T., Scorzetti G., Fell J.W., Kurtzman C.P., 2011. *Candida* Berkhout (1923) *In* The Yeasts, a Taxonomic Study, 5th edn, Vol. 2, ed. by Kurtzman, C.P., Fell, J.W. and Boekhout, T., Elsevier, London, pp. 987–1278.

Leandro, M.J., Cabral, S., Prista, C., Loureiro-Dias, M., Sychrová, H., 2014. The high-capacity specific fructose facilitator ZrFfz1 is essential for the fructophilic behavior of *Zygosaccharomyces rouxii* CBS 732^⊤^ Eukaryot Cell 13, 1371–1379.

Leandro, M.J., Sychrova, H., Prista, C., Loureiro-Dias, M.C., 2011. The osmotolerant fructophilic yeast *Zygosaccharomyces rouxii* employs two plasma-membrane fructose uptake systems belonging to a new family of yeast sugar transporters. Microbiology 157, 601–608.

Leandro, M.J., Sychrova, H., Prista, C., Loureiro-Dias, M.C., 2013. ZrFsy1, a high-affinity fructose/H+ symporter from fructophilic yeast *Zygosaccharomyces rouxii*. PLoS One 8,e68165.

Libkind D., Hittinger C.T., Valério E., Gonqalves C., Diver J., Johnston M., Gonçalves P., Sampaio J.P., 2011. Microbe domestication and the identification of the wild genetic stock of lager-brewing yeast. Proceedings of the National Academy of Sciences 108, 14539–14544.

Lopandic, K., Pfliegler, W.P., Tiefenbrunner, W., Gangl, H., Sipiczki, M., Sterflinger, K., 2016. Genotypic and phenotypic evolution of yeast interspecies hybrids during high-sugar fermentation. Appl. Microbiol. Biotechnol. 100, 6331–6343.

Lopandic K., Tiefenbrunner W., Gangl H., Mandl K., Berger S., Leitner G., Abd-Ellah G.A., Querol A., Gardner R.C., Sterflinger K., Prillinger H., 2008. Molecular profiling of yeasts isolated during spontaneous fermentations of Austrian wines. FEMS Yeast Res 8, 1063–1075.

Lopandic, K., Zelger, S., Bánszky L.K., Eliskases-Lechner, F. and Prillinger, H., 2006. Identification of yeasts associated with milk products using traditional and molecular techniques. Food Microbiol. 23, 341–350.

McGovern, P.E., Zhang, J., Tang, J., Zhang, Z., Hall, G., Moreau, R.A., Nunez, A., Butrym, E.D., Richards, M.P., Wang, C-s., Cheng, G., Zhao, Z., 2004. Fermented beverages of pre- and proto-historic China. PNAS 101(51), 17593–17598.

McGovern, P.E., Hall, G.R., Mirzoian, A., 2013. A biomolecular archaeological approach to ‘Nordic Grog’. Danish Journal of Archaeology, 2(2), 112–131.

Mendes-Ferreira A., Cosme F., Barbosa C., Falco V., Ines A., Mendes-Faia A., 2010. optimisation of honey-must preparation and alcoholic fermentation by *Saccharomyces cerevisiae* for mead production. Int. J. Food Microbiol. 144, 193–198.

Noble, A.C., Bursick, G.F., 1984. The contribution of glycerol to perceived viscosity and sweetness in white wine. Am. J. Enol. Vitic. 35,110–112.

OIV Compendium of International Methods of Analysis of Wines and Musts Vol. 1 & 2, 2014 (http://www.oiv.int/oiv/info/enmethodesinternationalesvin?lang=en).

Oliveira B.M., Barrio E., Querol A., Perez-Torrado R., 2014. Enhanced enzymatic activity of glycerol-3-phosphate dehydrogenase from cryophilic *Saccharomyces kudriavzevii*. PLoS One 9:e87290.

Paget, C., Schwartz, J., Delneri, D., 2014. Environmental system biology of cold-tolerant phenotype in *Saccharomyces* species adapted to grow at different temperatures. Mol. Ecol. 23, 5241–5257.

Paoli, M., Anesi, A., Antolini, R., Geulla, G., Vallortigara, G., Haase, A., 2016. Differential Odour Coding Isotopomeres in the Honeybee Brain. Scientific Reports, 6:21893, 1-9, DOI:10:1038/srep21893.

Pereira, A.P., Mendes-Ferreira, A., oliveira, J.M., Estevinho, L.M., Mendes-Faia, A., 2013. High-cell-density fermentation of *Saccharomyces cerevisiae* for the optimisation of mead production. Food Microbiology 33, 114–123.

Pereira, A.P., Dias, T., Andrade, J., Ramalhosa, E., Estavinho, L.M., 2009. Mead production: Selection and characterization assays of *Saccharomyces cerevisiae* strains. Food Chem. Toxicol. 47, 2057–2063.

Pérez-Torrado, R., González, S.S., Combina, M., Barrio, E., Querol, A., 2015. Molecular and enological characterization of a natural *Saccharomyces uvarum* and *Saccharomyces cerevisiae* hybrid. Int. J. Food Microbiol. 204, 101–110.

Peynaud, E., Domerq, S., 1955. Sur les espèces de levures fermentant selectivement le fructose. Annales de l’Institut Pasteur 89, 346–351.

Pretorius I.S., 2000, Tailoring wine yeast for the new millennium: novel approaches to the ancient art of winemaking. Yeast 16, 675–729.

Rainieri S., Pretorius I.S., 2000. Selection and improvement of wine yeasts. Annals in Microbiology 50, 15–31.

Redzepovic S., Orlic S., Sikora S., Majdak a., kozina B., Volschenk H., Viljoen-Bloom M., 2003. Differential malic acid degradation by selected strains of *Saccharomyces* during alcoholic fermentation. Int J Food Microbiol 83, 49–61.

Rodrigues de Sousa H., Spencer-Martins I., Gonçalves P., 2004. Differential regulation by glucose and fructose of a gene encoding a specific fructose/H^+^ symporter in *Saccharomyces sensu stricto* yeasts. Yeast 21, 519–530.

Saberi, M., Seyed-allaei, H., 2016. Odorant receptors of *Drosophila* are sensitive to the molecular volume of odorants. Scientific Reports, 6:25103, 1-11, DOI:10.1038/srep25103.

Styger, G., Prior, P., Bauer, F.F., 2011. Wine flavour and aroma. J. Ind. Microbiol. Biotechnol. 38, 1145–1159. DOI 10.1007/s10295-011-1018-4.

Subden R.E., Krizus A., Osothsilp C., Viljoen M., Van Vuuren H.J.J., 1988. Mutational analysis of malate pathways in *Schizosaccharomyces pombe*. Food Research International 31, 37–42.

Swiegers, J.H., Bartowsky, E.J., Henschke P.A., Pretorius, I.S., 2005. Yeast and bacterial modulation of wine aroma and flavour. Australian Journal of Grape and Wine Research 11, 139–173.

Taing O., Taing K., 2007. Production of malic and succinic acids by sugar-tolerant yeast *Zygosaccharomyces rouxii*. Eur. Food Res. Technol. 224, 343–347.

Volschenk H., Viljoen M., Grobler J., Petzold B., Bauer F., Subden R.E., Young R.A., Lonvaud A., Denayrolles M., van Vuuren H.J.J., 1997. Engineering pathways for malate degradation in *Saccharomyces cerevisiae*. Nature Biotechnology 15, 253–257.

Volschenk H., van Vuuren H.J.J., Viljoen-Bloom M., 2003. Malo-ethanolic fermentation in *Saccharomyces* and *Schizosaccharomyces*. Current Genetics 43, 379–391.

